# Scratching in Style: 3D Printers as Plotters for Automated and Complex Wound Healing Assays

**DOI:** 10.1101/2024.09.19.613824

**Authors:** Hanjo Köppe, Magnus G. Richert, Debora Singer, Jorn Köppe, Mattes Köppe, Mladen Tzvetkov, Henry W. S. Schroeder, Sander Bekeschus, Sandra Bien-Möller

## Abstract

Scratch wound healing assays remain one of the most commonly used 2D migration assays to obtain a broad overview of the migration behavior of cultured cells. They are easy to perform and cost-effective but not standardized in terms of the geometric dimensions of the scratch, e.g., the scratch width and straightness of the line, especially when performed manually. Furthermore, conventional scratch assays only include a single scratch. This leads to the evaluation of a restricted cell population within the culture plates but not of all cells grown in the chamber. There are commercially available ’scratch-makers’ that are highly expensive and lack advanced functions such as scratching complex patterns. However, such complex scratch formations would make it possible to assess the entire cell population in a chamber and evaluate the scratch’s impact on, for example, migration-associated protein expression. To address this, we developed a novel method utilizing 3D printers as 2D plotters to allow repeatable scratch dimensions for varying pipette tips. This open-source and cost-effective system can be performed with any plate format from any manufacturer under culture hood conditions. The 2D plotter-based method we developed and validated provides highly reproducible and consistent scratch parameters for *in vitro* migration analyses.

## Introduction

Scratch wound healing assays (scratch assays) are, alongside cell exclusion methods, the most frequently used 2D migration assays for evaluating the migration behavior of cells *in vitro*. For this, cells are cultured until they reach confluence and then separated by pulling a pipette tip through the monolayer to create a gap (scratch) with two facing wound edges (1,2). This is usually performed manually, leading to a highly inconsistent wound morphology and making it difficult to compare between different experiments. Ensuring equal wound diameters is particularly important for quantifying cell migration behavior after different treatments, as these are regularly compared with a control group and depend on equal scratches in the control and intervention groups (8). Nevertheless, scratch assays remain widely used, with around 12,000 PubMed entries, most of which are performed manually. Due to the natural limits of the human hand, manual execution inevitably leads to inconsistencies and irregularities in form, contact pressure and pulling speed. The conventional method also lacks the opportunity of a scalable approach, as it is almost impossible to scratch manually within the 96-well plate format. Moreover, complex shapes cannot be implemented manually, although this would allow migration processes to be induced in cells from various regions of a well. This would make it possible to investigate the regulation of migration within the cell assembly and draw conclusions about the regulation of migration-associated proteins, e.g., by immunofluorescence staining after the experiment. Such a versatile use of scratch assays underlines the importance of making standardized approaches accessible to the majority, thus improving various fields of science.

Although different methods exist to create equal gaps for migration analyses, none fully meets the requirements regarding cost efficiency, reliability, user-friendliness, and scalability (9). For example, cell exclusion assays (5), in which cells are seeded into silicon chambers so that a barrier separates them until the chambers are removed to create a gap, are highly versatile but have some limitations. The current compartment-based approaches are not scalable, as the chambers do not fit into smaller plate formats and are only available in certain geometric dimensions and volumes. In addition, depending on the desired experimental observation, it may also be that the surrounding cells are not optimally affected to exactly mimic a ’scratch assay’. This could be important when simulating, for example, surgical injuries or the formation of scars from repeated scratching (10).

There are also some highly functional DIY approaches (3,6,7,11), but most are over-engineered and require sophisticated machinery to produce, making them non-reproducible for most scientists. Recently, *Lin et al.* (12) published an approach to use a programmable, open-source drawing robot to create scratches for studying migration behavior. Although this approach also involves low production effort, parts still need to be 3D printed for the setup, which means the opportunity to use the 3D printer as a more direct solution has been missed so far. Furthermore, most DIY machines are tailored to one specific size of cell culture plates and only fulfill a particular need. Some companies also offer commercially available devices that can perform single straight gaps, but these are extremely expensive (around USD 12,000) and thus not suitable for everyone. *Fenu et al.* (13) published an interesting low-cost approach that showed that it is possible to reproduce single straight lines with high precision by sliding magnets through the wells. Still, again, it lacks the same capabilities as the cell exclusion inserts. Notably, there are some other methods to perform scratch assays with high accuracy, e.g., thermal ablation of cells within the gap by laser radiation (14,15) or electrical current (16), but all of them have their disadvantages regarding user-friendliness, effectiveness, and low-cost. Consequently, there is a vast need for standardization and automation of the conventional pipette-based scratch assay through an open hardware and software approach.

All of the above-mentioned challenges that the other approaches fail to address can be solved using the precise XYZ axis environment provided by 3D printers. Linear- guided axes and stepper motors eliminate hand tremor and imprecise tip movements. Spring-loaded test tips ensure constant contact, vector graphics enable free-form scratches, and an open-source program for generating G-code allows adjustments to meet individual preferences. We have named our approach **ASAPR**, which stands for **A**dvanced **S**cratch **A**ssay **P**lotting **R**obot. ASAPR offers some significant benefits: 1) any culture plate or culture dish can be used (up to a certain size), 2) any pipette tip can be used, 3) any 2D shape can be scratched, 4) all device parts with cell contact can be sterilized, 5) the procedure is carried out under a sterile culture hood, 6) the method offers automatic washing steps to avoid cross-contamination 7), the system is programmable to the desired needs 8), it is easy to set up, and 9) requires only minor extra costs (∼15 USD) besides the 3D printer itself.

## Results

### Setup and principle of ASAPR on 3D printers

ASAPR can be used with any commercially available 3D printer that supports the execution of ’raw’ or ’manual’ G-code, which is the case for most models. The idea is to use the 3D printer as a plotter by moving the print head only in the x and y direction while giving the z-axis a fixed execution height within the desired culture dish or plate. We have therefore developed two different approaches for holding the pipette. The first uses a pipette mount which is screwed directly in the M6 thread of the printhead as a replacement for the nozzle. The second approach uses a 3D printed custom attachment for a precision chuck which then holds the pipette tip (Fig. 1A). We recommend the first approach, as it is easier to determine the required X and Y offsets later (see Methods) and the automatic leveling of the 3D printer is more effective as the pipette tip is in exactly the same position as the nozzle. To ensure uniform pressure throughout the experiment, we used high-precision spring-loaded test probes, which are typically used for ’in-circuit tests’. The end of the tip itself consists of a regular pipette tip, which is cut by a 3D-printed cutting device to ensure a uniform length and then placed on the tip of the spring-loaded test probe. The pipette tip centers itself due to the conical shape of the spring-loaded probe tip. To hold the cell culture plates on the printing platform, we designed a 3D-printed mount that clamps downwards (data available on GitHub). This mount can be adjusted in size via an adjusting screw to allow the plates to be clamped tightly. Other mounts, e.g., for dishes, will be released in the future on GitHub. At this point, the mechanical setup is already complete, and the first G-code can be generated using the provided program ’ASAPR’ (Fig. 1B, for setting up the option to include washing steps, see GitHub).

**Figure 1.**
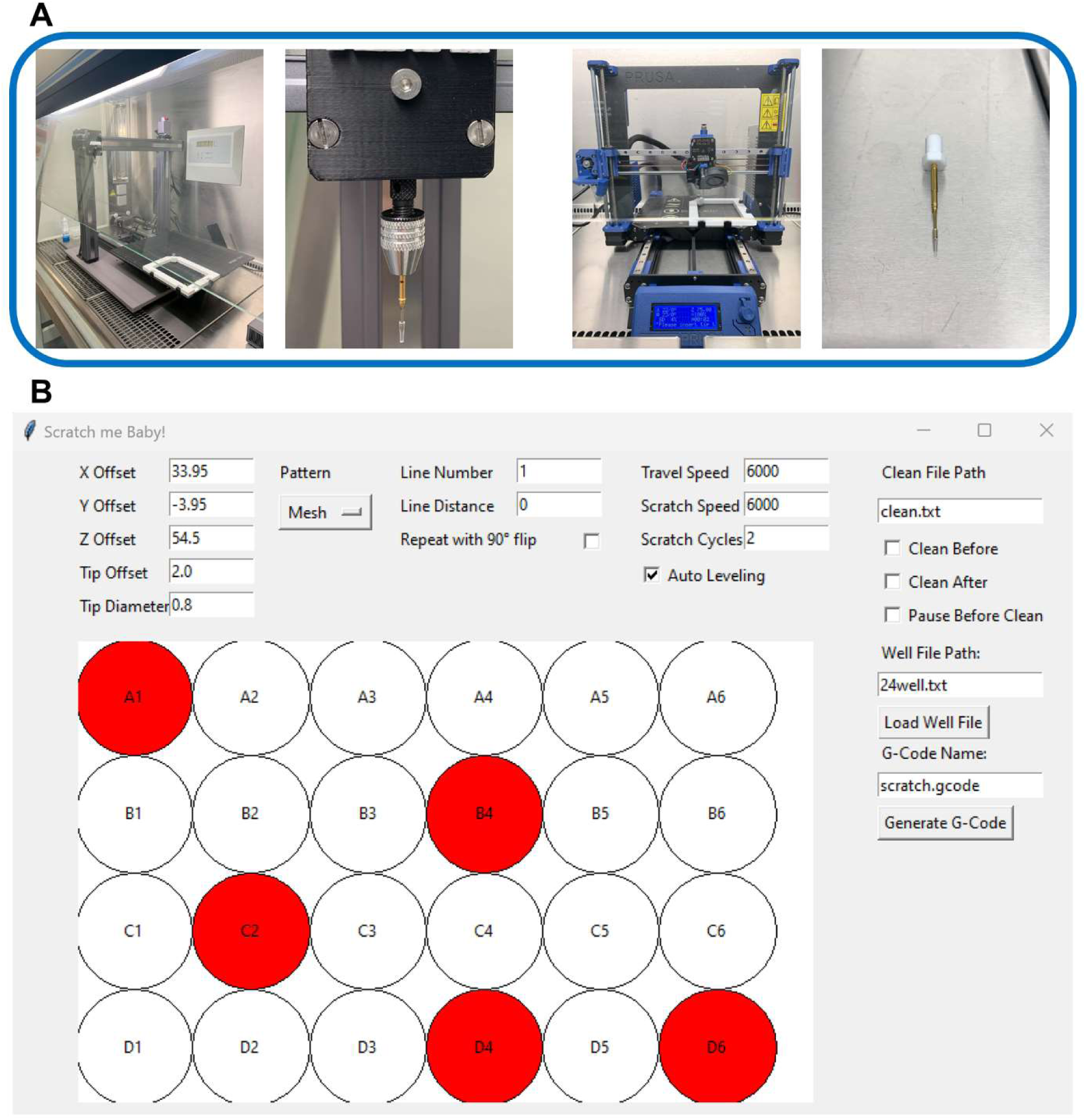
Setup of ASAPR on 3D printers and G-code generating program. (A) Overview image of the two printers and the mounting options used. On the left, the Snapmaker A350 with the custom-made mount (II), and on the right, the Prusa MK3S+ with linear rails and the 3D-printed M6 holder (I) can be seen. Also visible is the 3D-printed plate holder that is clamped onto the printing platform, which enables the cell culture plates to be clamped firmly in the same position. (B) An example representation for the generation of the G-code for individual straight scratches within a 24-well plate. A safety distance of 2 mm to the edges of the well is set via the ’Tip Offset’ task function. The running and scratch speeds are set to 6000 mm/min. The scratch movement runs back and forth to achieve the straightest possible wound edges, which is set via 2 ’Scratch Cycles’. Wells of the culture plate that are not required (here A1, B4, C2, D4, and D6) were omitted by clicking on them and are then not included in the code. After generation, the code is saved in the same folder as the graphical user interface (GUI).

### Generation of appropriate G-code

The print head does not automatically know at which section of the printing platform the cell culture plate is located. For this purpose, the X- and Y-axis offsets must first be determined to specify the maximum frame within which the print head may later operate. The z-axis offset must also be determined so that the tip acts at the correct depth, although this does not have to be as precise as in the X and Y directions, as the spring-loaded tip compensates for unevenness in conjunction with the printer’s (auto)-leveling function. It is highly recommended that the offsets be carefully determined, as they must be defined once, and they will always remain the same for the device afterward (for full calibration, see GitHub). G-code works similarly to a series of directional instructions in a three-dimensional coordinate system. We have written a Python program ’ASAPR’ to generate these instructions. This program provides a variety of functions and offers three different patterns. The *Mesh* pattern only executes straight lines, which can be placed at any distance from each other and can also be used for individual scratches. These lines can also be tilted by 90 degrees to create a grid or cross. The *Circle* pattern is used to scratch circles that increase in size into larger cavities to encourage all cells in a well to migrate. The *Scalable Vector Graphics* (*SVG)* pattern makes it possible to scratch any conceivable .svg file, i.e., vector graphics, by automatically scaling them to the size of the wells. Vector graphics can be created beforehand using open-source software such as Inkscape. The G-code can then be transferred to the 3D printer as specified in the instruction manual of the 3D printer. The geometric dimensions of the plate to be processed can be taken from the data sheet and are loaded into the program via a text file, which allows the use of many plate types and quick adjustments to the orientation by swapping X and Y (the exact options are discussed on GitHub).

### Validation of single-scratch reproducibility

We first validated the reproducibility of standard single straight lines, as primarily performed using conventional scratch assays. Three 24-well plates were processed, so 72 scratches were included for the single scratch validation. The scratching experiments were carried out on the glioblastoma cell line LN-18. The optimum conditions before scratching were determined to be almost 100% confluence, short incubation times to minimize cell-cell adhesion and cell-surface adhesion, scratching speeds of 100 mm/s or higher and the use of two ’Scratching Cycles’. To ensure reproducibility, we focused on the parameters wound width and wound area, whereby the standard deviation of the wound width can also be used to assess the straightness of the lines. Our approach resulted in a coefficient of variation of 1.61% for wound area and wound width (Fig. 2A) with a perfect correlation (r = 0.99, data not shown) between each other, indicating a rectangular shape. Furthermore, no significant difference in wound width was found between the columns or the rows of the three 24-well plates (Fig. 2C). A comparison with self-performed manual scratching does not appear expedient, as the subjective bias would be too significant, which is why only published data were used for comparison. Based on the published data on the accuracy of manually performed scratch assays, the coefficient of variation is approximately 8-17% for the gap width and about 6% for the straightness of the line (13,17).

**Figure 2.**
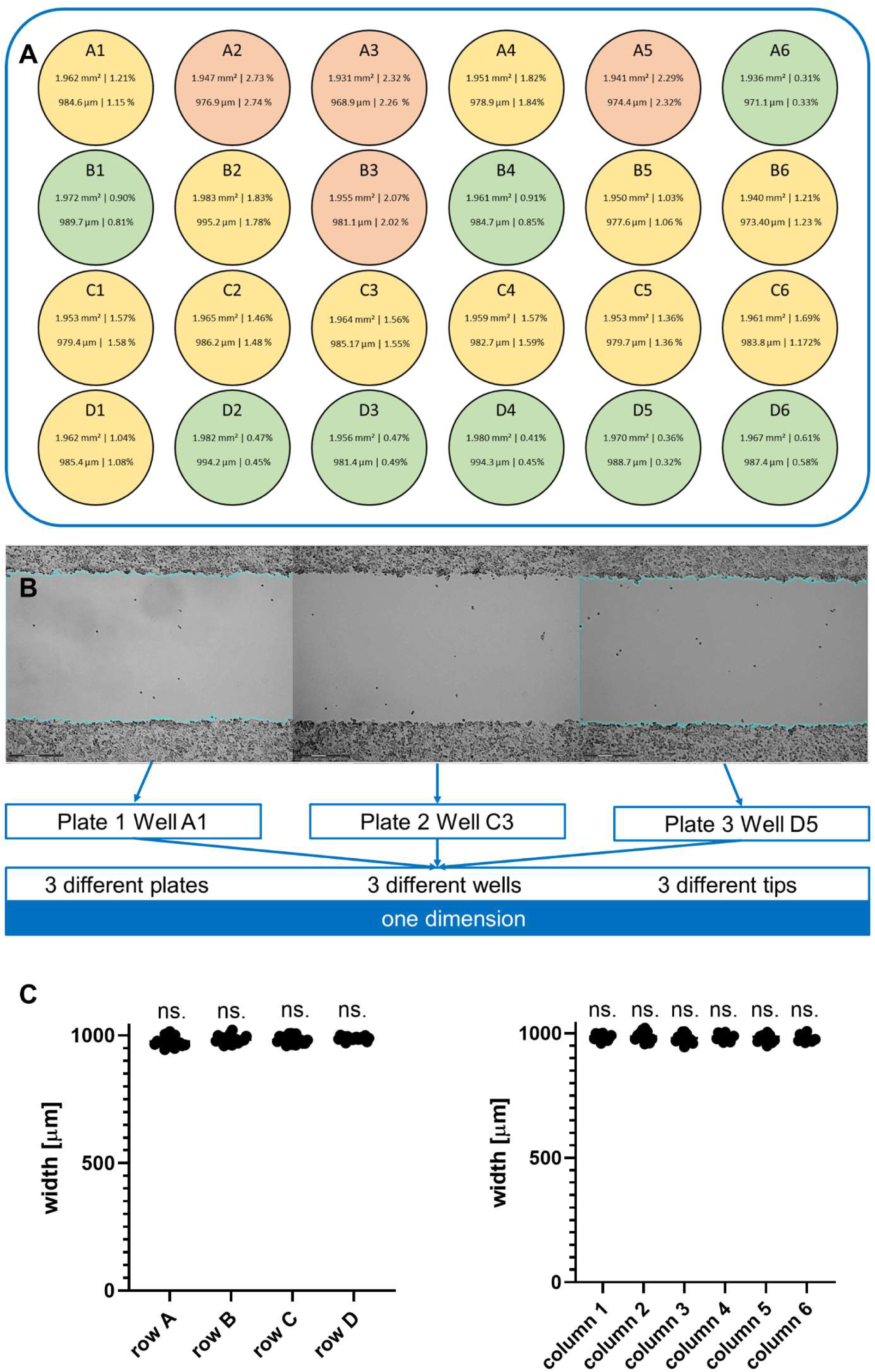
Single Scratch validation. (A) The heatmap of the individual cell culture wells shows the cell-free area (top) and the wound width (bottom) as well as the corresponding coefficients of variance (CV, in %), demonstrating the consistency in making single straight scratches in 24-well plates cultured with the glioblastoma cell line LN-18 using ASAPR. The area is given in mm² and the width in µm. Green wells represent a CV of less than 1%, yellow wells have a CV of 1-2%, and red wells show a CV of more than 2% for the mean gap area and mean gap width, as they correlate well. All scratches were generated using the modular 3D printer *Snapmaker A350* with a 200 µl pipette mounted. Each image was taken randomly across the scratches on the first plate, and the position was then marked to be viewed on the other two plates as well. No shift in the localization was detected in the other scratches. (B) Representation of three example scratches from three different wells of three different plates, all created with ASAPR. The images were taken using brightfield microscopy. Scale bar: 300 µm. (C) A comparison between all rows and columns also showed no significant differences in the wound width generated. Mean + single measurement points. Rows n = 18; columns n = 12. Statistical significance was calculated by one-way ANOVA with Tukey’s multiple comparisons test. ns. = non significant.

### Evaluation of differences between different printers

To investigate whether the geometric dimensions of scratches differ when using different 3D printers, we examined the processing of a 24-well plate between two printers. For this purpose, we compared the scratching results from the previously utilized *Snapmaker A350* with the *Prusa MK3S+* that had been upgraded with linear rails (Fig. 3A). Two different mounting methods for the spring-loaded testing tip were used: our preferred method of screwing in a 3D-printed M6 thread (I) (Fig. 1A, right) and a custom-made mount (II) (Fig. 1A, left). Both approaches produced highly reproducible results. As expected, there were slight differences between the two printers due to machine-specific vibrations and stepper motor speeds. The *Prusa MK3S+* produced straighter scratches with a standard deviation of only 14.13 µm across the entire scratch width. The *Snapmaker A350* performed slightly worse, with a standard deviation of 18.29 µm. This difference was within the expected range, as its significantly larger print area likely resulted in less rigid axes. Due to the presumably lower vibrations and faster printing speed of the *Prusa MK3S+*, the overall scratch width was also slightly smaller, 925.7 µm on average, compared to 971.2 µm with the *Snapmaker A350* (Fig. 3B). However, the difference in average width was only 4.68%, suggesting that while it may be beneficial to standardize on a single printer for internal consistency, this does not challenge the underlying concept of the method itself. These results further support the assumption that axis speed positively correlates with the straightness of the scratches, as it likely increases the inertia of the tip and minimizes vibrations of the rails.

**Figure 3.**
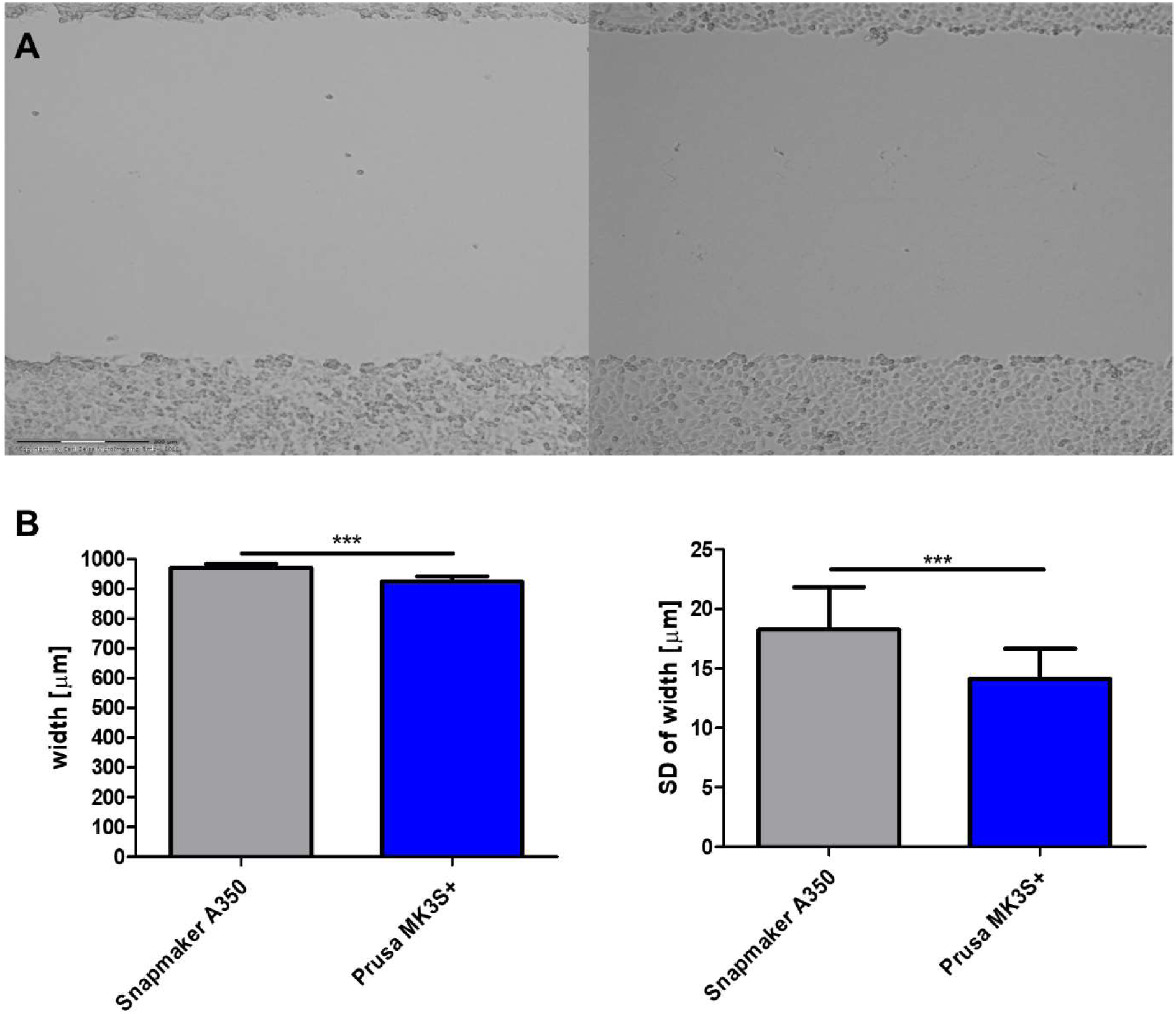
Comparison of scratching results between different printers. (A) A sample scratch from the Snapmaker A350 (left) and the Prusa MK3S+ (right) demonstrated the minimal variation in width. Scale bar: 300 µm. (B) Differences in wound width and standard deviation of wound width between the two different printers used. The deviations are highly significant, primarily due to the very low intra-assay variability. Mean ± SD, n = 24. Statistical significance was calculated by unpaired, two-tailed t-test. *p ≤ 0.05, ** p ≤ 0.01, *** p ≤ 0.001.

### Validation of complex two-dimensional shapes

The study of complex scratch patterns is particularly interesting because it allows for the investigation of the influence of scratch morphology itself on migration behavior (18). Additionally, when combined with live cell imaging, it becomes possible to observe a small section and the entire well, hopefully leading to more precise overall results. It is impossible to perform manually (21). The G-code generating program creates a toolpath similar to a laser engraver and follows it. As a result, the smallest possible details also depend on the pipette tip size used. The tip widths are approximately 1000 µl - 1.2 mm, 200 µl - 0.8 mm, and 10 µl - 0.7 mm. However, this does not reflect the expected scratch width, as the pipette tip vibrates within the well and pushes a cluster of cells ahead of it, so the actual scratch width is typically larger. To validate the scratching of complex two-dimensional shapes, mazes were applied in 6-well plates (Fig. 4A), which were subsequently quantified by fluorescence microscopy. For this, LN-18 cells were stained with Vybrant DiD Cell-Labeling Solution 24 hours before the scratch was done. The quantification of the cell-free area resulted in a mean area of 4.91×10^8^ µm^2^ and a coefficient of variation of 1.51% for a total of 18 wells with respect to the wound area. Quantifying shape accuracy is challenging as it would require an approach that overlays all shapes and determines the deviations from the original shape. Furthermore, three additional complex .svg files were converted into G-code using our program (Fig. 5). The first file was created in Inkscape to demonstrate compatibility between the two programs. The other two graphics were selected to better illustrate the range of possibilities and to show that even highly complex shapes can be realized.

**Figure 4.**
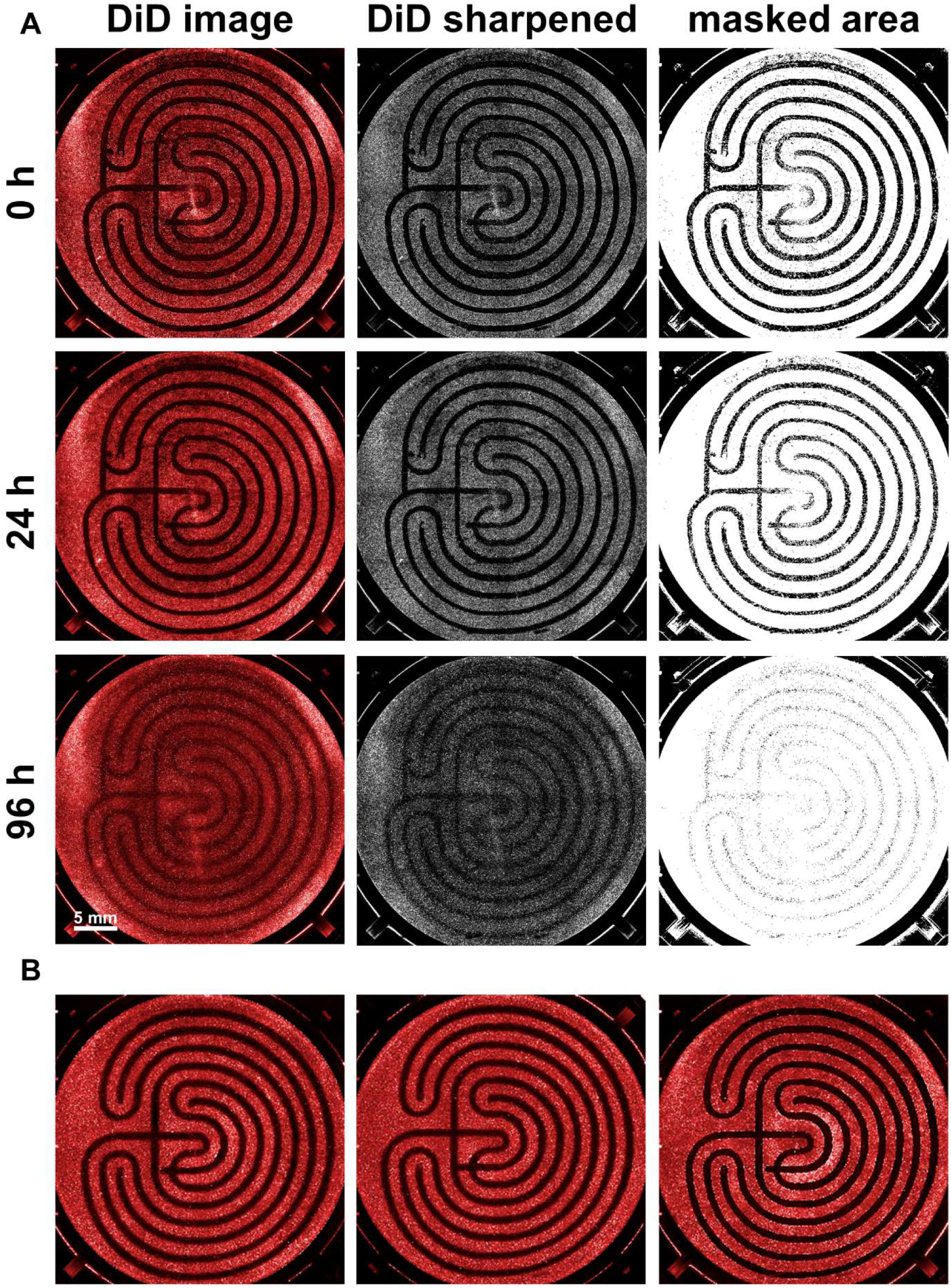
Quantification of complex shapes, e.g., mazes. (A) The image processing procedure is shown after the immunofluorescence images have been recorded by scanning all wells of a 6-well plate. The image is first increased in contrast to enhance the distinguishability, and then only the non-covered area is masked. Over 96 hours, it can be seen how the wound area closes almost completely and uniformly. Hydroxyurea was added as a proliferation inhibitor. Scale bar: 5 mm. (B) Three further example images of scratched mazes at time point t=0h demonstrate the high replicability. Cells were stained with Vybrant DiD Cell-Labeling Solution.

**Figure 5.**
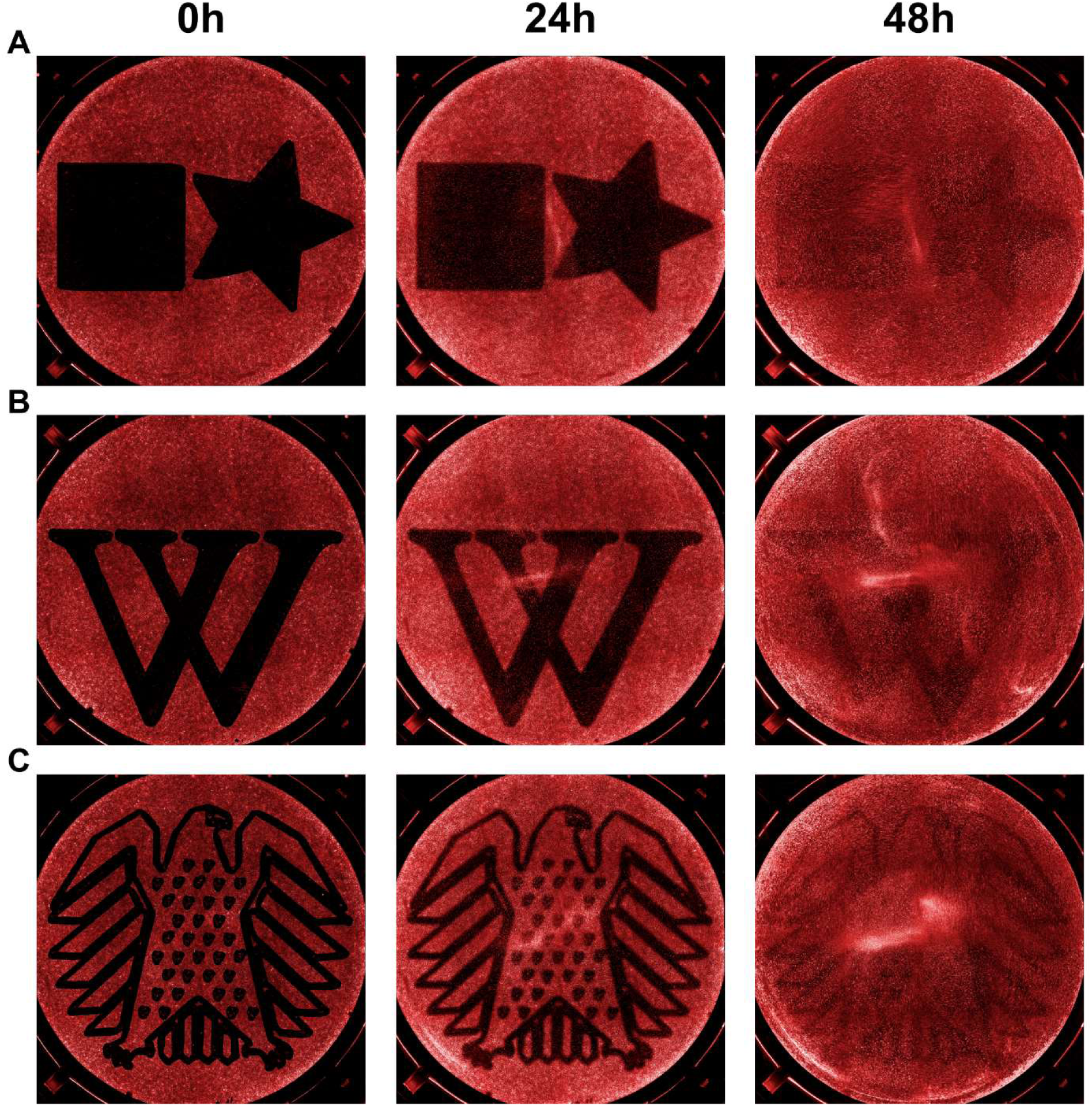
Examples of complex vector graphics. (A) The star and square were created using the open source software Inkscape and were imaged for 48 hours. In these experiments, no hydroxyurea was added as a proliferation inhibitor so the closure can be attributed more to proliferation. (B) The "W" logo from Wikipedia and (C) the German Bundestag coat of arms are shown. The wells of the 6-well plates have a diameter of 34.6mm. Cells were stained with Vybrant DiD Cell-Labeling Solution.

### Induction of migration-associated proteins and cytokines

To stimulate almost all cells in a 6-well plate to migrate, the wells were treated with a total of 16 individual scratches across the entire width, all equally spaced, resulting in 18 cell bridges of 400 µm width from which the cells migrate left and right to cover the gap (Fig. 6A). While in single scratches only cells near the wound edges migrate, this approach achieves a migration status in almost all cells in response to this large wound area, allowing for subsequent analysis of regulatory mechanism linked to cell damage and migration in most cells. This approach is called the ’Mesh’. After 24 hours of migration, the medium was collected to determine IL-6 secretion by ELISA. IL-6 signalling promotes a variety of activities that support gliomagenesis including cell invasion and migration and was therefore picked as an example cytokine (28). Additionally, cells were either harvested for subsequent Western blot analysis or stained with crystal violet to quantify the relative cell number by staining all viable cells.

**Figure 6.**
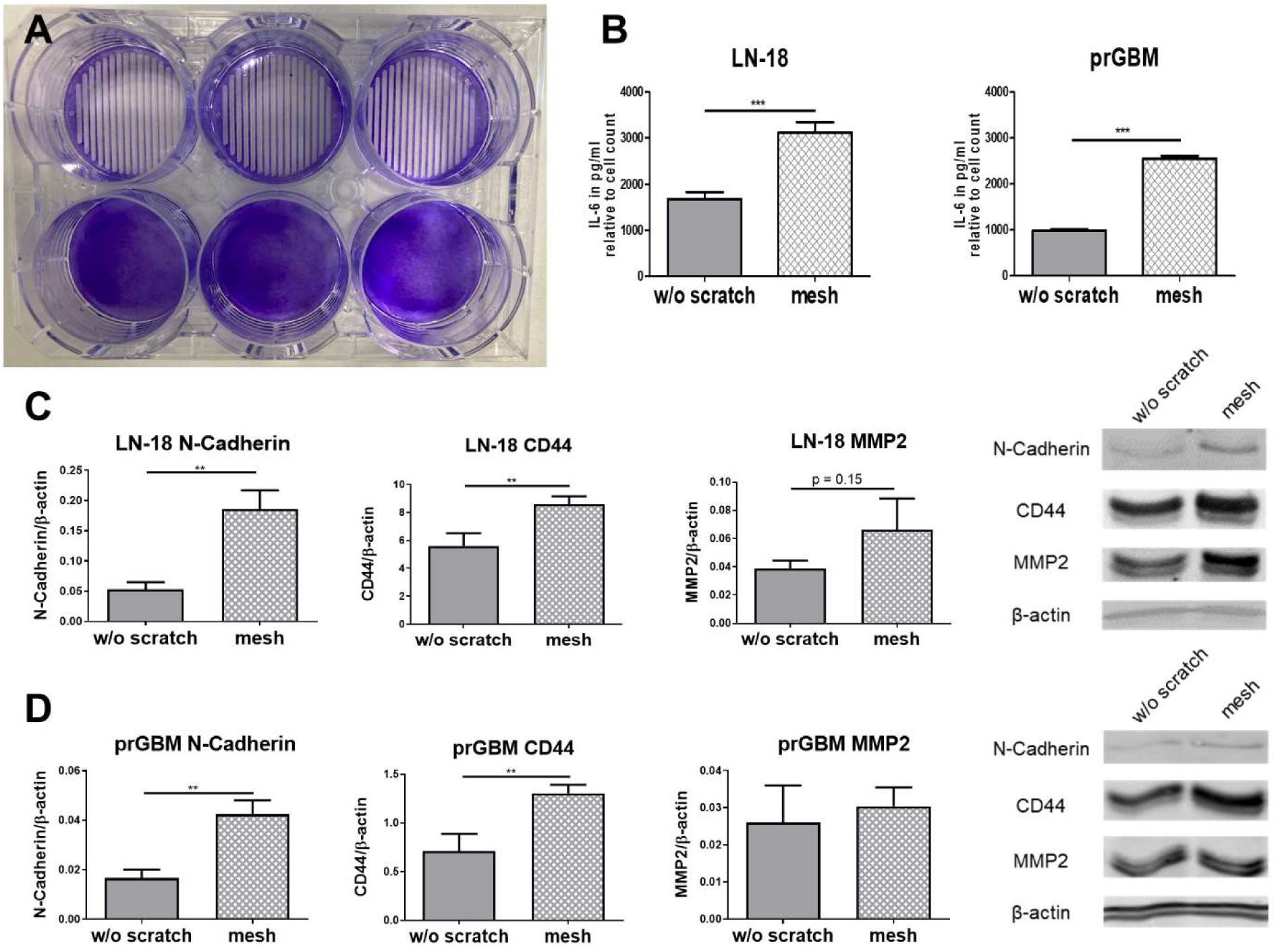
Modulation of migration-related proteins and secretion of IL-6 by the scratching procedure in glioblastoma cells. (A) Cells were grown to full confluence and kept untreated (bottom) or mesh-scratched (top) prior to crystal violet staining (LN-18 cells). Visible are the 400 µm wide cell bridges, which are evenly arranged across the entire well. (B) To investigate whether migrating cells secrete more IL-6 than non-migrating cells, the cell monolayer of a 6-well plate was scratched over the entire well after the cells had reached confluency using the mesh procedure. The IL-6 concentrations are determined by ELISA and normalized to the relative cell number within a well by staining all viable cells with crystal violet after removing the cell culture supernatant. All conditions were treated with 5mM Hydroxyurea during migration period to inhibit proliferation. Mean ± SD, n = 3, unpaired two-tailed t-test, *p ≤ 0.05, ** p ≤ 0.01, *** p ≤ 0.001. (C) After 24h of migration, the cells were harvested, and protein was isolated for subsequent Western blot analysis of the migration-related proteins CD44, MMP2 (Matrix-Metalloproteinase 2), and N-Cadherin. Mean ± SD, n = 3, unpaired two-tailed t-test *p ≤ 0.05, ** p ≤ 0.01, *** p ≤ 0.001.

**Figure 7.**
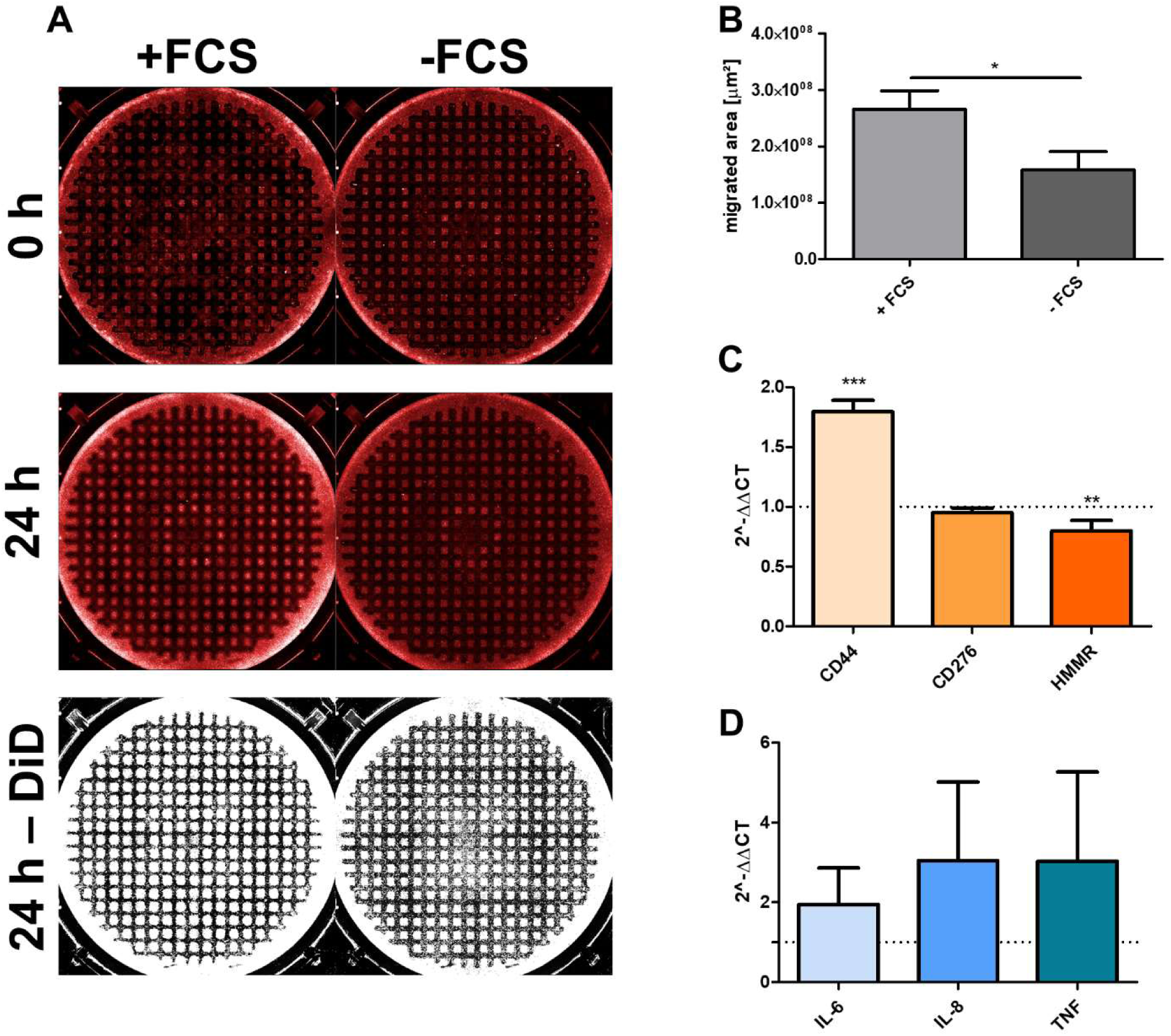
Differential migration profile under varying culture conditions. (A) An example experiment in which different migration rates were analyzed under FCS-containing and FCS-free culture conditions during migration period in LN-18 cells is shown. After 24 hours, it is visible that the cell region stained with Vybrant DiD Cell-Labeling Solution is significantly larger in the FCS-containing medium. (B) The mean migration area within 24 hours was about 1.68 times larger in the FCS-containing cell culture compared to the FCS-free culture (0.266 cm^2^ vs. 0.158 cm^2^). All conditions were treated with 5mM Hydroxyurea during migration period to inhibit proliferation. Mean ± SD, n = 3, unpaired two-tailed t-test, *p ≤ 0.05, ** p ≤ 0.01, *** p ≤ 0.001. (C-D) Relative mRNA expression of the migration-related proteins CD44, CD276, and HMMR (C) and the cytokines IL-6, IL-8, and TNFα (D) in LN-18 glioblastoma cells in FCS-containing and FCS-free culture conditions after 24 hours of migration. The FCS-free experiments were used as controls, with the relative expression for each mRNA set to 1. Mean ± SD, n = 4, one-way ANOVA with Dunnett post-test, *p ≤ 0.05, ** p ≤ 0.01, *** p ≤ 0.001.

This allowed us to compare protein and cytokine levels between migrating cells from scratched wells and non-migrating cells from those without scratches. In addition to LN-18 cells, IL-6 secretion and expression of selected migration-associated proteins (CD44, N-Cadherin, and MMP2) (30,31,32) were also examined in primary glioblastoma cells (prGBM) in the same experimental setting. The cells from the scratched wells showed a significant induction of CD44 and N-Cadherin for both LN- 18 and prGBM after 24h of migration. The protein content of matrix metalloproteinase- 2 (MMP2) was also elevated 1.7-fold in LN-18 cells, but this did not reach statistical significance, while MMP2 was not altered in prGBM cells. The strongest induction was observed for N-cadherin in LN-18 cells, with a 3.5-fold increase after 24 hours. CD44 protein expression was increased 1.5-fold and 1.8-fold in scratched LN-18 and prGBM cells after 24 hours, respectively. Additionally, the amount of IL-6 secreted per cell doubled in LN-18 and increased 2.5-fold in prGBM cells 24 hours after the scratch. It is important to note that all cells were treated with 5 mM hydroxyurea after scratching to determine only migration and not proliferation. As the aim here is merely to demonstrate a possible application and not to analyze which migration proteins are regulated in glioblastoma cells through migratory processes, the results will not be further explained at this point.

### In-depth migration analyses

Conventional scratch assays are poorly suited for investigating the influence of potential migration inhibitors on protein and mRNA levels after migration has occurred. If, for example, the cells were subsequently lysed to analyze the modulation of proteins involved in migration, e.g., using Western blot or RT-PCR analysis, only the cells that form the wound gap would have to be isolated, as only they have migrated. Otherwise, the remaining cells within a well might overshadow or at least diminish the observed effect. To accurately track the influence of different conditions on migration, we developed the mesh further by adding the option of a 90° rotation, thus creating a grid. This results in hundreds of small cell islands of approximately the same size, which migrate concentrically and independently of each other. This setup allows for a more precise analysis of how different conditions affect cell migration, as all cell islands migrate simultaneously. Moreover, there is increasing evidence that scratch assays only provide relatively accurate results if the entire wound area is considered and not just a part of the scratch region (26). In this context, our approach, combined with live-cell imaging, significantly improves the experimental setup, as it allows for observing the migration behavior of almost all cells within a given culture condition.

In a proof-of-concept experiment, we compared the migration rates and the expression of migration-related markers in FCS-free cultures versus cultures supplemented with 10 % FCS for 24h as both conditions are considered to be common positive and negative controls (29). As almost all cells within the well are under migration conditions, this setup allows for directly assessing how different culture conditions influence migration behavior. The migration markers were extended beyond CD44 and IL-6 to include the second hyaluronic acid receptor HMMR (33), CD276 (34), which is involved in the epithelial-mesenchymal transition, as well as the two cytokines IL-8 (35) and TNF (36), both of which play important roles in glioblastoma. Since this is merely an experimental example setup, the connections between the markers and the rationale for their selection are not discussed in detail at this point. Contrary to expectations, not all investigated migration markers were upregulated by the increased energy availability provided by FCS. Specifically, CD44 (mean = 1.79) was upregulated as expected, CD276 was not differentially expressed at all (mean = 0.95), and HMMR (mean = 0.80) was even slightly downregulated at the mRNA level. The cytokines IL- 6, IL-8, and TNFα showed a general trend towards higher mRNA expression, but the variation was too large to be statistically significant.

This experimental setup also makes it possible to investigate the influence of the scratch itself on the cells and the subsequent activation of signaling cascades. For example, cells could be analyzed at different time points to better understand the temporal dynamics of specific migration cascades. Furthermore, placing a new grid over the previous one after wound closure would be conceivable to investigate potential anticipatory effects associated with faster secondary wound closure.

## Discussion

We have established and validated a 3D printer-based method that provides highly reproducible and consistent scratch parameters for *in vitro* migration analyses. However, during the test phase, numerous sources of error emerged that should be considered when using ASAPR. Many of the adjustments/adaptations made are also crucial for improving conventional scratch assays. The area to be scratched should be considered as an elastic sheet that adheres to a surface. If the cells are too tightly connected, more neighboring cells are pulled out than just those that should be lost by scratching with the pipette tip, resulting in frayed wound edges (19). However, if the cells adhere too strongly to the plate, the pipette tip may not catch all of them, resulting in very irregular wound edges (Supplementary Figure S1). There are two possible solutions to eliminate these two interferences, both of which should ideally be implemented simultaneously. First, the cells should be given a short adhesion time (optimally no longer than 24 hours) to minimize too close cell-cell and cell-surface contacts (19). Second, the scratch should be made at very high speeds to take advantage of the inertia of the cells adhering to the surface. Overall, optimal conditions in our experimental setting were almost 100% confluence, short incubation times to minimize cell-cell adhesion and cell-surface adhesion, scratching speeds of 100 mm/s or higher, and using two ’scratching cycles’.

Additionally, the inertia of the pipette tip increases at higher speeds, which should theoretically lead to less vibration and a high straightness of the scratches. In this context, it would be exciting to see what significantly faster printers could achieve, as the *Snapmaker A350* used here is limited to a theoretical maximum of 130 mm/s due to its lead screw mechanism. Although the Prusa MK3S+ is equipped with linear rails and has higher maximum speeds (theoretical up to 250mm/s), the limited range within a well means that acceleration is the most critical factor. Core-XY and rail-guided printers, in particular, can reach much higher speeds and accelerations (22).

A further significant improvement was achieved through the use of spring-loaded test probes. Although most 3D printers have a well-functioning automatic leveling system, it was still common that the cell surface was not reached with the same z-value in certain places. As the pipette tips themselves are very inflexible, this is an adjustment that is difficult to achieve in manually performed scratch assays, and it cannot be guaranteed that the same pressure is applied consistently to the entire surface. This incertitude in manual scratching was eliminated by using spring-loaded test probes. Although our approach has yielded excellent results, some experimental areas can still be improved. For example, while the spring-loaded test probes have greatly improved the consistency of the pressure applied, their slight but present backlash can lead to irregularities that result in less-than-optimal outcomes which gives room for further improvement.

The exact XYZ-axis environment of 3D printers, which forms the basis of the ASAPR concept, proves to be an ideal framework for using scratch assays. It enables much more precise, reproducible, and customizable assay configurations than manually performed scratch assays (13, 17). Moreover, this approach opens up a whole new world of possibilities for 2D migration analyses, as the induction of migration by complex scratch patterns enables, for the first time, a simple comparison between migrating and non-migrating cells as well as the analysis of the scratch’s impact. The study of complex scratching patterns is fascinating because it allows the influence of scratching morphology on migration behavior to be investigated. While in single scratches, only cells near the wound edges migrate, this approach achieves a migration status in almost all cells in response to this large wound area, allowing subsequent analysis of the regulatory mechanisms associated with cell damage and migration. Our preliminary experiments using the mesh configuration in glioblastoma cells showed regulation of selected migratory-relevant proteins such as CD44 and N-Cadherin by Western Blot analysis. In addition, immunofluorescence microscopy and live cell imaging are also possible as part of these experiments and might show expression differences directly in the individual migrated cells.

However, this open-source platform offers many future opportunities, particularly regarding software development and component improvement. The time saved by ASAPR is also worth mentioning, as the scratches are not only made faster and more precisely but also consistently in the same location. This is particularly beneficial when capturing images manually, without a live-cell imager, making the process more convenient because the images can always be taken at the same position. This precise reproducibility, even in terms of spatial resolution, offers new approaches for migration studies. For example, repeated scratches could be applied to the same site, enabling the investigation of pathomechanisms in scar formation or multiple wound damages, as well as subsequent analyses of signaling cascades and proteins involved in the migration processes.

Overall, ASAPR represents an efficient approach requiring minimal production effort, as all components can be manufactured using the 3D printer. Of note, many DIY approaches to perform highly reproducible scratch assays require production-based 3D printing, each with strengths and weaknesses (23,24). The approach by *Lin et al.* (12) is particularly noteworthy as it comes closest to our concept. However, as 3D- printed parts were needed without using the printer, a straightforward process was overlooked. With 3D printers becoming more common in many laboratories for various reasons (25), the barrier to entry for ASAPR is relatively low, facilitating the performance of reproducible, precise, and standardized scratch assays with the opportunity for more in-depth migration analyses.

## Conclusion

The presented method will allow more significant variation in 2D cell migration experiments. It is also economical to implement and open-source. In the future, setups combining 3D printing and variations thereof may explore versatile migration analysis in all three dimensions.

## Acknowledgments

The authors thank Tina Sonnenberger (Department of Pharmacology, Centre of Drug Absorption and Transport (C_DAT), University Medicine of Greifswald, Greifswald, Germany) for her excellent technical assistance.

## Author Contributions

Conceptualization, H.K., M.G.R., J.K. and M.K.; methodology, H.K., M.G.R., J.K. and M.K.; coding, M.G.R and M.K.; CAD modelling, J.K.; validation, H.K.; formal analysis, H.K., S.B.-M., S.B. and D.S.; investigation, H.K. and D.S.; resources, H.W.S.S., M.V.T. and S.B.; data curation, H.K. and S.B.-M.; writing—original draft preparation, H.K.; writing—review and editing, H.K., S.B.-M., S.B., D.S., M.V.T. and H.W.S.S.; supervision, S.B.-M., S.B., M.V.T. and H.W.S.S.; project administration, S.B. and S.B.-M.; funding acquisition, M.V.T., H.W.S.S., S.B.-M., S.B. All authors have read and agreed to the published version of the manuscript.

## Funding statement

High-content imaging was accomplished by funding from the German Federal Ministry of Education and Research (BMBF), grant number 03Z22Di1 (to S.B.).

## Data availability

All printing components, technical instructions, and operating principle and setup explanations are available on the project’s GitHub page (https://github.com/MagnusRichert/ASAPR) and from the corresponding author upon request.

## Conflicts of Interest

The authors declare no conflict of interest.

## Methods

### Cell culture

The human glioblastoma cell line LN-18 (ATCC, Manassas, VA, USA) was used to develop and validate 3D printer-based scratch assays. Further, primary glioblastoma (prGBM) cells from a surgically resected patient sample were used as described before (20). GBM cells were cultured at 37 °C, 95 % relative humidity, and 5 % carbon dioxide fumigation in DMEM supplemented with 10% FCS, 2 mM glutamine, and non-essential amino acids. All cell lines were routinely monitored for mycoplasma contamination using a PCR-based assay. The cells were plated at a density of 1.2 × 10^6^ cells per well in 6-well format or 0.1 × 10^6^ cells per well in 24-well format plates and incubated until confluence was reached.

### ASAPR hardware setup

A detailed explanation of the operating principle of ASAPR and instruction for the 3D printed parts, along with a comprehensive calibration and setup guide, can be found on GitHub. Below is a brief explanation of the approach used for the *Snapmaker A350* (SNAPMAKER HK, Hong Kong, China). A custom holder was printed, which was mounted directly onto the X-axis with four M6 screws so that the system can operate without a print head. A precision drill chuck (MBS03; Donau Elektronik, Metten, Germany) was inserted into this holder and secured with a set screw. The drill chuck held a spring-loaded test probe (2021-B-1.5N-NI-0.8; PTR HARTMANN, Werne, Germany). A 200 μl pipette tip was used, which was cut to the same length using a 3D-printed device. This cut-off tip was attached to the test probe and centers itself due to its conical shape. The optimal spring travel was set at 1 mm. The 3D-printed plate frame was attached with four clamps designed for glass print beds. This frame includes an M3 knurled screw for securely clamping the inserted plates.

The *Prusa MK3S+* (Prusa Research, Prag, Czech Republic) was purchased second-hand via a marketplace, already upgraded with linear rails on the X- (https://www.printables.com/model/16838-prusa-mk3-mk3s-mk3s-x-axis-linear-rail-guide-upgra) and Y-axes (https://www.printables.com/model/5620-prusa-mk3-mk3s-mk3s-y-axis-linear-rail-guide-upgra). An M6 thread was printed to secure the spring-loaded probe tip, into which a precision probe sleeve (PTR H 1015 L, PTR HARTMANN, Werne, Germany) was inserted. This holder was screwed into the M6 mount of the print head until it reached the stop after each homing procedure. The jerk setting in the system settings had to be limited to a maximum of 15mm/s, as the printer would occasionally skip steps otherwise. The same frame as for the *Snapmaker A350* was used to clamp the cell culture plates.

### Scratch configuration and post-scratch processing

The scratches for single straight lines were made using the specified parameters in the G-code generating program: tip offset ’2’[mm], mode ’Mesh’, line number ’1’, line distance ’0’, travel speed ’6000’[mm/min], scratching speed ’6000’[mm/min], auto- leveling ’on’, scratch cycles’ 2’. The scratches for the mesh were made with the specified parameters in the G-code generating program: tip offset ’2’[mm], mode ’Mesh’, line number ’18’, line distance ’1.6’[mm], travel speed ’6000’[mm/min], scratching speed ’6000’[mm/min], auto-leveling ’on’, scratch cycles’ 2’. The scratches for the .svg files were made using the specified parameters in the G-code generating program: tip offset ‘2’[mm], mode ‘svg’, svg scale ‘100’[%], travel speed ‘6000’[mm/min], scratching speed ‘6000’[mm/min], auto-leveling ‘on’, scratch cycles ‘1’. The svg files are displayed on GitHub. After scratching, the culture medium was removed, and the cells were washed three times with PBS to remove detached cells and prevent their reattachment. For migration experiments over 24h, pre-warmed serum free culture medium or medium containing 10 % FCS and 5 mM hydroxyurea as a proliferation inhibitor was added. For disinfection, the tip was immersed in 70% ethanol for atleast 30s before and after scratching and then allowed to air dry. Contamination has never occurred during testing period.

### Scratch microscopic and evaluation

Single straight wounds were imaged using the PALM Robo software of the AxioVision HXP 120C microscope with a 5× objective and a 10× ocular (Carl Zeiss Microscopy, Jena, Germany), and the exact position of the scratches was saved to analyze the same area after incubation. Fluorescence images were captured using a high-content imaging device (PerkinElmer, Hamburg, Germany). For this, the cells were stained with Vybrant DiD Cell-Labeling Solution (V22887; Thermo Fisher Scientific, Darmstadt, Germany) 24h before imaging. FIJI (https://imagej.net/software/fiji) was used to process all images in terms of contrast and brightness and to calculate the wound areas for the scratch assays using the wound healing plugin of Suarez-Arnedo et al. (4). The plugin settings remained the same for evaluation of all images: variance window radius: 3; threshold value: 30; percentage of saturated pixels: 0.001; set scale global?: yes; the scratch is diagonal?: yes. The analysis of fluorescence images was performed using *Harmony 4.9* software (PerkinElmer, Hamburg, Germany).

### Crystal violet staining

The cell culture supernatant was aspirated from scratched and control cells, and adherent cells were rinsed once with PBS. Afterward, cells were fixed in 4 % paraformaldehyde for 10 min, then gently rinsed thrice with PBS. 300 μl of a 0.5 % crystal violet staining solution (Sigma-Aldrich, Deisenhofen, Germany) was added to the cells, and the plates were incubated at room temperature for 10 min. The cells were washed several times with distilled water until the dye stopped coming off. After washing, the plate was carefully tapped on filter paper to remove any remaining liquid. Finally, the stained cells were treated with 300 µL SDS solution (1 %) and incubated on a shaker for 10 min to dissolve the dye. The optical density was determined at 560 nm (OD560) using a multiplate reader and 36 flashes per well (Tecan Infinite M200, Crailsheim, Germany).

### Western Blotting

GBM cells were scraped, transferred to a 1.5 mL tube, and centrifuged at 10,000 rpm for 3 min. Afterward, the cell pellets were resuspended in lysis buffer (50 mM Tris-HCl, 100 mM NaCl, 0.1% Triton X-100, and 5 mM EDTA, pH 7.4) containing 1 mM leupeptin, 1mM PMSF and 1 mM aprotinin and incubated on ice for 30 min, followed by centrifugation at 12,000 rpm for 5 min at 4 °C. The supernatant was used for protein determination using the BCA method. Subsequently, after denaturation in Laemmli SDS sample buffer at 90 °C for 10 min, 30 μg of protein was separated on 10% SDS polyacrylamide gels. The tank blot system (Bio-Rad, Hempstead, UK) was used for immunoblotting of the separated proteins onto Whatman nitrocellulose membranes, which were then blocked for 1 h at room temperature in 5 % FCS in Tris-buffered saline containing 0.05 % Tween 20 (TBST). The membrane was incubated overnight at 4 °C under rotation with the following primary antibodies diluted in TBST (1:1000): mouse anti-CD44 (Cell Signaling Technology, Boston, MA, USA); rabbit anti-MMP2 (Cell Signaling Technology, Boston, MA, USA); mouse anti-N-Cadherin (Cell Signaling Technology, Boston, MA, USA) and mouse anti- β -actin (Santa Cruz Biotechnology, Inc., Heidelberg, Germany). Afterward, the membrane was rinsed three times with TBST for five minutes each time, followed by incubation with the secondary fluorescence-labeled antibody for one hour at room temperature with gentle shaking: anti-mouse IRDye 680 or 800 CW or anti-rabbit IRDye 680 or 800 CW (all from Li-cor Bioscience, Bad Homburg, Germany). After incubation, the membrane was rinsed three times with TBST for five minutes at room temperature on a horizontal shaker. Fluorescence signals were detected using the Odyssey CLx Imaging System (Li-cor Bioscience, Bad Homburg, Germany). The relative fluorescence intensities of the specific bands were calculated and normalized to β-actin as a loading control.

### Real-Time PCR

Total RNA was isolated using TRIzol (Thermo Fisher Scientific, Darmstadt, Germany) together with RNeasy Kit (Qiagen, Hilden, Germany) and reversely transcribed using the High-Capacity cDNA Reverse Transcription Kit (Applied Biosystems, Weiterstadt, Germany). The following Gene Expression Assays on Demand (Applied Biosystems) were used: IL-6, Hs00985639_m1; IL-8, Hs00174103_m1; TNF, Hs01113624_g1; HMMR, Hs00234864_m1; CD276, Hs00987207_m1; CD44, Hs01075864_m1 endogenous control eukaryotic TBP, Hs00427620_m1; GAPDH, Hs99999905_m1; and B2M, Hs99999907_m1.

Quantitative Real-Time PCR was performed on a QuantStudio 7 Flex Real-Time PCR System from Applied Biosystems. All measurements were taken as technical triplicates. The mRNA content of target genes was normalized to the geometric mean of TBP, B2m, and GAPDH and is expressed relative to the control cells (1 = 100%) as described before (27).

### ELISA for measurement of IL-6

The supernatant of the cultured GBM cells was transferred to a 1.5 ml tube, centrifuged at 1,000 rpm for 3 min, and pipetted into a new 1.5 ml tube to remove debris. Afterward, the samples were stored at -80°C until measurement. Before measurement, the samples were diluted 1:10 to stay within the calibration curve. The concentrations of IL-6 in the supernatants were determined using ELISA kits (D6050; R&D Systems, Minneapolis, Minnesota, USA) according to the manufacturer’s instructions.

### Statistical analysis

Statistical analyses were performed using the *prism* 5.0 (GraphPad Software, San Diego, CA, USA). All data are presented as mean ± SD of at least three independent experiments. Pairwise comparisons were performed using Student’s t-test and one-way ANOVA with Dunnett post-test or Tukey’s multiple comparisons test for multiple groups. Statistical significances were defined as * p < 0.05; ** p < 0.01; and *** p < 0.001.

---

**Figure S1.**
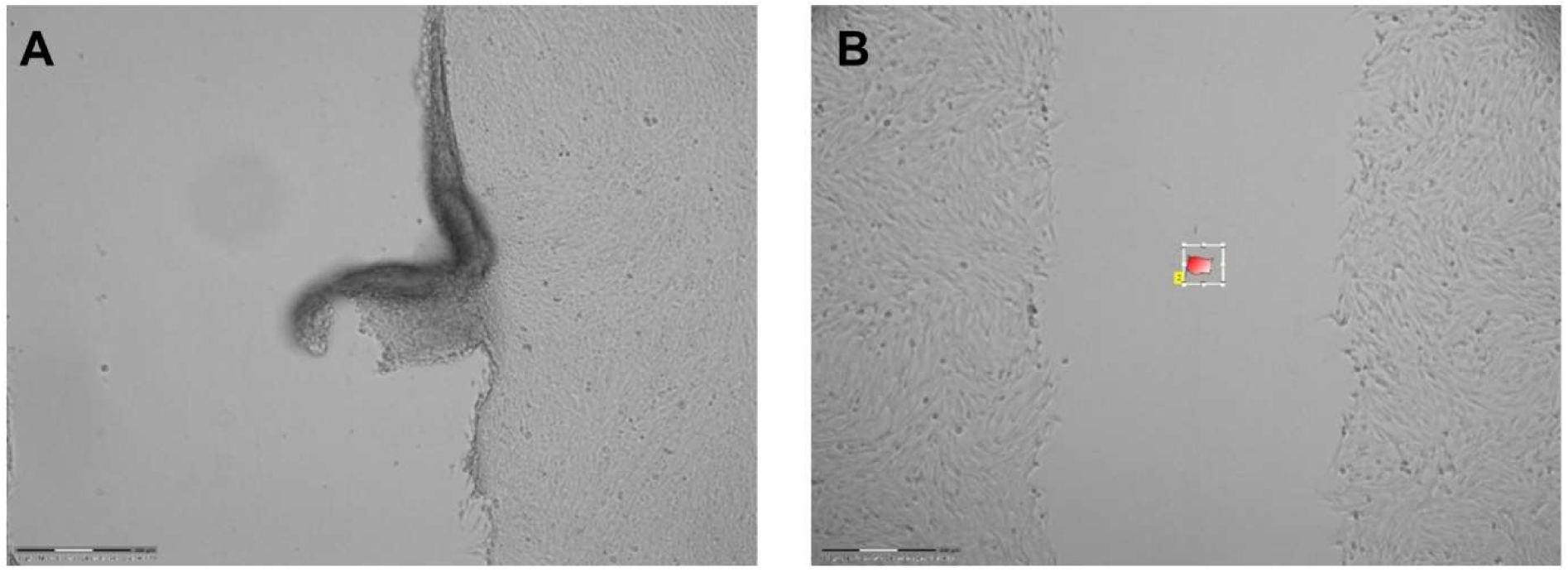
Examples of irregular scratches. (A) Shown is a failed scratch due to firm cell-cell contacts. It is visible how an entire layer of cells has detached while maintaining contact with the wound edge. (B) Visible irregular wound edges as a consequence of robust cell- surface contacts. Scale bar: 300 µm.

## Notes

https://github.com/MagnusRichert/ASAPR

### Competing Interest Statement

The authors have declared no competing interest.

### Summary of Updates

Title updated; Figure 1 updated and rearranged; Figure 2 updated and supplemented by row and column comparison; Figure 3 added for printer comparison; Figure 5 added to show dealing with vector graphics; Figure 7 added to give outlook and utilisation example; Updated formatting; Methods clearified; Conclusion, Acknowledgments, Author contributions and Funding statement added; References added; Fixing of general spelling mistakes;

## References

1. Liang CC, Park AY, Guan JL. In vitro scratch assay: a convenient and inexpensive method for analysis of cell migration in vitro. Nat Protoc. 2007;2(2):329–333.

2. Martinotti S, Ranzato E. Scratch Wound Healing Assay. Methods Mol Biol. 2020;2109:225–229.

3. Gough W, Hulkower KI, Lynch R, et al. A quantitative, facile, and high-throughput image-based cell migration method is a robust alternative to the scratch assay. J Biomol Screen. 2011;16(2):155–163.

4. Suarez-Arnedo A, Torres Figueroa F, Clavijo C, Arbeláez P, Cruz JC, Muñoz-Camargo C. An image J plugin for the high throughput image analysis of in vitro scratch wound healing assays. PLoS One. 2020;15(7):e0232565.

5. Shiode Y, Morizane Y, Matoba R, et al. A novel cell exclusion zone assay with a barrier made from room temperature vulcanizing silicone rubber. PLoS One. 2017;12(12):e0190198.

6. Chen S, Bin Abdul Rahim AA, Mok P, Liu D. An effective device to enable consistent scratches for in vitro scratch assays. BMC Biotechnol. 2023;23(1):32.

7. Grimmig R, Babczyk P, Gillemot P, Schmitz K-P, Schulze M, Tobiasch E. Development and Evaluation of a Prototype Scratch Apparatus for Wound Assays Adjustable to Different Forces and Substrates. Applied Sciences. 2019; 9(20):4414.

8. Nyegaard S, Christensen B, Rasmussen JT. An optimized method for accurate quantification of cell migration using human small intestine cells. Metab Eng Commun. 2016;3:76–83.

9. Radstake WE, Gautam K, Van Rompay C, et al. Comparison of in vitro scratch wound assay experimental procedures. Biochem Biophys Rep. 2023;33:101423.

10. Nikolić DL, Boettiger AN, Bar-Sagi D, Carbeck JD, Shvartsman SY. Role of boundary conditions in an experimental model of epithelial wound healing. Am J Physiol Cell Physiol. 2006;291(1):C68–C75.

11. Acosta S, Canclini L, Galarraga C, Justet C, Alem D. Lab-made 3D printed stoppers as high-throughput cell migration screening tool. SLAS Technol. 2022;27(1):39–43.

12. Lin, Y., Silverman-Dultz, A., Bailey, M., & Cohen, D. J. (2024). A programmable, open-source robot that scratches cultured tissues to investigate cell migration, healing, and tissue sculpting. Cell reports methods, 4(12), 100915.

13. Fenu M, Bettermann T, Vogl C, et al. A novel magnet-based scratch method for standardisation of wound-healing assays. Sci Rep. 2019;9(1):12625.

14. Mathioudaki E, Rallis M, Politopoulos K, Alexandratou E. Photobiomodulation and Wound Healing: Low-Level Laser Therapy at 661 nm in a Scratch Assay Keratinocyte Model. Ann Biomed Eng. 2024;52(2):376–385.

15. Wu SY, Sun YS, Cheng KC, Lo KY. A Wound-Healing Assay Based on Ultraviolet Light Ablation. SLAS Technol. 2017;22(1):36–43.

16. Keese CR, Wegener J, Walker SR, Giaever I. Electrical wound-healing assay for cells in vitro. Proc Natl Acad Sci U S A. 2004;101(6):1554–1559.

17. Jin W, Shah ET, Penington CJ, McCue SW, Chopin LK, Simpson MJ. Reproducibility of scratch assays is affected by the initial degree of confluence: Experiments, modelling and model selection. J Theor Biol. 2016;390:136–145.

18. Vedula SR, Leong MC, Lai TL, et al. Emerging modes of collective cell migration induced by geometrical constraints. Proc Natl Acad Sci U S A. 2012;109(32):12974–12979.

19. Harris AR, Peter L, Bellis J, Baum B, Kabla AJ, Charras GT. Characterizing the mechanics of cultured cell monolayers. Proc Natl Acad Sci U S A. 2012;109(41):16449–16454.

20. Marx S, Wilken F, Wagner I, et al. GD2 targeting by dinutuximab beta is a promising immunotherapeutic approach against malignant glioma. J Neurooncol. 2020;147(3):577–585.

21. Jonkman JE, Cathcart JA, Xu F, et al. An introduction to the wound healing assay using live-cell microscopy. Cell Adh Migr. 2014;8(5):440–451.

22. Wolf J, Werkle KT, Möhring HC. Study on Dynamic Behaviour in FFF 3D-printing with Crossed Gantry Kinematic. Procedia CIRP. Volume 121. 2024. 162-167.

23. Joshi AS, Madhusudanan M, Mijakovic I. 3D printed inserts for reproducible high throughput screening of cell migration. Front Cell Dev Biol. 2023; 11:1256250. Published 2023 Aug 30.

24. Acosta S, Canclini L, Galarraga, et al. Lab-made 3D printed stoppers as high-throughput cell migration screening tool. SLAS Technology. 2022;27(1):39–43

25. Saggiomo V. A 3D Printer in the Lab: Not Only a Toy. Adv Sci (Weinh). 2022;9(27):e2202610.

26. Balko, S., Kerr, E., Buchel, E., Logsetty, S., & Raouf, A. (2023). A Robust and Standardized Approach to Quantify Wound Closure Using the Scratch Assay. Methods and protocols, 6(5), 87.

27. Vandesompele J, De Preter K, Pattyn F, et al. Accurate normalization of real- time quantitative RT-PCR data by geometric averaging of multiple internal control genes. Genome Biol. 2002;3(7)

28. West AJ, Tsui V, Stylli SS, et al. The role of interleukin-6-STAT3 signalling in glioblastoma. Oncol Lett. 2018;16(4):4095–4104.

29. Justus CR, Leffler N, Ruiz-Echevarria M, Yang LV. In vitro cell migration and invasion assays. J Vis Exp. 2014;(88):51046. Published 2014 Jun 1.

30. Noronha C, Ribeiro AS, Taipa R, et al. Cadherin Expression and EMT: A Focus on Gliomas. Biomedicines. 2021;9(10):1328. Published 2021 Sep 26.

31. Inoue A, Ohnishi T, Nishikawa M, et al. A Narrative Review on CD44’s Role in Glioblastoma Invasion, Proliferation, and Tumor Recurrence. Cancers (Basel). 2023;15(19):4898. Published 2023 Oct 9.

32. Du R, Petritsch C, Lu K, et al. Matrix metalloproteinase-2 regulates vascular patterning and growth affecting tumor cell survival and invasion in GBM. Neuro Oncol. 2008;10(3):254–264.

33. Pibuel MA, Poodts D, Molinari Y, et al. The importance of RHAMM in the normal brain and gliomas: physiological and pathological roles. Br J Cancer. 2023;128(1):12–20.

34. Liu S, Liang J, Liu Z, et al. The Role of CD276 in Cancers. Front Oncol. 2021;11:654684. Published 2021 Mar 26.

35. Sharma I, Singh A, Siraj F, Saxena S. IL-8/CXCR1/2 signalling promotes tumor cell proliferation, invasion and vascular mimicry in glioblastoma. J Biomed Sci. 2018;25(1):62. Published 2018 Aug 8.

36. Ramaswamy P, Goswami K, Dalavaikodihalli Nanjaiah N, Srinivas D, Prasad C. TNF-α mediated MEK-ERK signaling in invasion with putative network involving NF-κB and STAT-6: a new perspective in glioma. Cell Biol Int. 2019;43(11):1257–1266.

